# Head and gaze tracking of unrestrained marmosets

**DOI:** 10.1101/079566

**Authors:** Hjalmar K. Turesson, Thamiris Botelho Ribeiro Conceição, Sidarta Ribeiro

**Affiliations:** York University, Toronto, Canada; Universidade Federal do Rio Grande do Norte, Natal, Brazil; Instituto do Cérebro, Universidade Federal do Rio Grande do Norte, Natal, Brazil

## Abstract

New technologies for manipulating and recording the nervous system allow us to perform unprecedented experiments. However, the influence of our experimental manipulations on psychological processes must be inferred from their effects on behavior. Today, quantifying behavior has become the bottleneck for large-scale, high-throughput, experiments. The method presented here addresses this issue by using deep learning algorithms for video-based animal tracking. Here we describe a reliable automatic method for tracking head position and orientation from simple video recordings of the common marmoset (*Callithrix jacchus*). This method for measuring marmoset behavior allows for the estimation of gaze within foveal error, and can easily be adapted to a wide variety of similar tasks in biomedical research. In particular, the method has great potential for the simultaneous tracking of multiple marmosets to quantify social behaviors.

## Introduction

Recent technological developments allow us to record [10,12,61] and manipulate [4,42,47] the nervous system with unprecedented precision and scale. Yet, the psychological relevance of our sophisticated manipulations and large-scale recordings can only be inferred from their effects on behavior. Today, properly quantifying behavior has become the main bottleneck for high-throughput experiments [7,56]. It is common practice to apply standard tests designed to measure psychological constructs such as anxiety and spatial memory, for example, by using the Elevated Plus Maze [41] or the Morris Water Maze [35]. Such testing requires animals to be individually handled, making data acquisition labor intensive, increasing costs and reducing experimental throughput. Alternatively, various simple detectors (e.g. capacitance sensors or photo-beams) can be arranged to automatically acquire data at specific sites (e.g. drinking [17] and feeding [14] stations). This type of automation allows for high throughput behavioral quantification, but fails to capture complex or subtle behaviors such as social interactions through gaze behaviors. A more promising approach is to record high-dimensional data from sensor arrays (e.g. video) and extract relevant information using computer vision algorithms. This approach has the potential to provide a better characterization of behavior, capable of automatically capturing complex and subtle behaviors, while simultaneously reducing both cost and labor intensity [46]. Raw video frames are composed of a high number of pixels whose values do not straightforwardly correlate with an animal’s behavior. In order to reliably measure behavior give such variability, many computer vision systems rely on explicitly-designed dimensionality reduction methods that extract specific features (e.g. to achieve illumination invariance [18]). The result of this preprocessing can then be used to measure or classify behavior [24,62]. However, the explicit design of feature extraction methods that are robust under a wide range of conditions is difficult and time-consuming [2]. Deep learning algorithms, in contrast, automatically learn a hierarchy of increasingly abstract representations, running from simple feature detectors (e.g. edge detectors) to the final classification. This strategy obviates the need for manually designed feature extraction and has proven successful in a great variety of tasks [2,30]. A particular group of deep learning algorithms, convolutional neural networks (CNNs), is inspired by the local connectivity pattern of the visual cortex [63], and are optimized for data organized in arrays like those produced by a digital camera. The recent development of CNNs have led to impressive improvements on visual recognition tasks [29], making them the most popular deep learning method in computer vision and ideal for video-based tracking of behavior.

Most work on the automatic measurement of behavior has been done on fruit flies [5,51–53], zebra fish [39,40] and mice [36,38]. However, despite the importance of non-human primates as research models in several areas of life sciences the methods development is lagging behind. In particular, the common marmoset *(Callithrix jacchus)* is becoming an increasingly important primate model in biomedical [21,37,45,54,64], genetic [23,25,44], and neuroscience [28,49] research. The growing popularity of marmosets stems from their similarity to humans regarding the disease susceptibility profile [11,55], their relative ease of handling, high fecundity and fast development [43], and the recent development of key tools for genetic manipulations [44,48] and neuroscience experiments [37]. To the best of our knowledge, only one publication has reported on a preliminary method for automated behavioral tracking of marmosets [6].

In contrast to rodents, primates in general have to orient their gaze precisely which makes gaze direction informative of what they pay attention to [33,49]. Further, a striking feature of the gaze behavior of small-headed primates such as marmosets is their rapid head movements. When shifting their gaze, marmosets tend to move their heads in quick jerks, similar to how bigger primates move their eyes when making saccades. Although marmosets can and do make saccadic eye movements, they are limited to the central 10° [33], with the result that head movements contribute substantially to the final gaze shift [20,32]. This might be explained by these primates’ small head and relatively large eyes [16,57]. The lesser head size results in lower rotational inertia which in turn, lowers the muscular force required to produce rapid head movements. Since rotational inertia decreases faster with size than muscle force does, head movements become gradually more favorable with smaller head sizes. Thus, the combination of a small fovea and a big contribution of head movement to the gaze direction makes head tracking of marmosets remarkably informative, allowing gaze estimation without eye measurements. For marmosets, head orientation alone indicates marmosets’ gaze direction within ±10°. By tracking a marmoset’s head position and direction, we get a rich source of information about how it moves and what captures its interest. We here present a video-based method that uses a CNN to track head position and orientation (thus approximating gaze direction) in the common marmoset.

## Results

We set out to develop a video-based method for automatic tracking of head position and gaze direction of marmosets. To this end, we first recorded marmosets engaged in a 15-minute vocal learning experiment using a video camera mounted above the test box (**Fig 1**). We annotated a subset of the video frames with head direction and trained a CNN on this data. Head direction was explicitly predicted by the model, whereas head position was indirectly estimated from the model’s spatial activation pattern [65].

**Fig 1.**
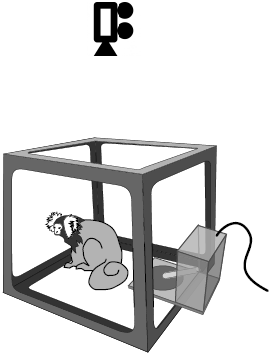
Experimental setup. Video was recorded by a camera placed above the subject while it was taking part in a vocal learning experiment. The device to the right of the subject is the reward dispenser with spout (light tube) and recipient below (dark disc).

There were two sources of uncertainty in the input data. First, humans do not score the video frames in an entirely consistent fashion, but will vary a bit from frame to frame in their estimate of identical head directions. Second, head direction angles (0 to 360°) were binned into 28 classes resulting in a loss of precision. To provide an estimate of the first source of uncertainty, two investigators annotated the same 1,361 frames and the differences between the estimated directions and positions were measured. Between the two investigators, the mean difference was 9.2° (median 7.2°) for direction, and 10 pixels (median 9 pixels) for position. We refer to these differences as the inter-human disagreement. Beyond measuring the uncertainty in the input data, the inter-human disagreement also provides a reference to which the model’s performance can be compared. We operationalized the model error as the difference between a human scorer’s estimate of head direction and position, and that of the model.

In order to address the limits of the input data and to contain the effects of over-fitting, we used model averaging [50,51]. Unless otherwise noted, the results presented below originate from the averaging of 30 models. **Fig 2A** shows an example of predicted head direction and position in comparison with the human annotation in a sequence of video frames. Note that although the manually marked position is shown, this is only to evaluate the model’s performance and no explicit position information was used in training the model.

**Fig 2.**
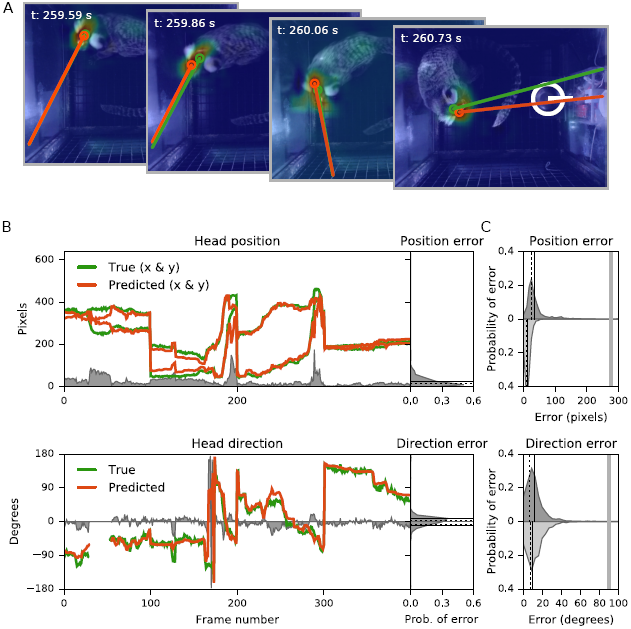
Head tracking. **A**, Four video frames illustrating the tracking of head direction and position. The time of each frame is shown on the upper left corner. The orange line and circle show the predicted head direction and position, respectively. The green line and circle show the corresponding manual annotations. The CAM (see Materials and Methods) is superimposed on the video frames as a heat map with more discriminative image regions shown in warmer colors. The white circle and horizontal bar in the leftmost frame show the location of the reward dispenser. **B**, Left panels show position (upper panel) and direction (lower panel) tracking of 400 frames of the test data. Orange lines show predicted head position coordinates in pixels (upper), and head direction in degrees (lower). Green lines show the manual annotation, and gray shows the model’s error. In the lower panel, the light green and red fields around frames 25 – 50 indicate agreement (green) and disagreement (red), about frames where the animal’s gaze was directed out of the horizontal plane. The right panels show histograms of the error from the left panels (left and right panels share y-axis). **C**, Position (upper) and direction (lower) errors across all test data. The model errors are shown in dark gray, and the corresponding inter-human disagreements are shown in light gray. Black lines show mean (full lines) and median (dotted) error, and the vertical gray lines indicate chance error. The x-axes have been truncated for visual clarity.

To systematically test tracking performance we annotated 2,000 video frames with head direction and position. This allowed us to compare the model’s predictions with the inter-human disagreement. Randomly selecting the frames from the training set is likely to overestimate performance since adjacent frames in these videos often are highly correlated. Therefore, to test the model, we used labeled frames from five videos (400 frames from each video) that did not contribute to neither training nor validation data. The five videos included all three subjects. **Fig 2B** shows the predicted head position and orientation, as well as the associated errors, in one of the five videos. The magnitude of the head direction error is reported degrees from 0 to 180° and the position error is reported in pixels (640 × 480-pixel video frames). As an upper limit for the errors they can be compared to chance, which is 90° for direction, and 277 pixels for position. For the data in **Fig 2B**, the mean direction error was 9.1° (median 3.3°), and the mean position error was 24 pixels (median 16 pixels).

**Fig 2C** shows the direction and position error on all the test data compared with the inter-human disagreement. The mean head direction error was 10.9° (median error 6.2°), which is very close to the inter-human disagreement with a mean of 9.2° (median 7.2°). The mean position error was 33 pixels (median 23 pixels), compared to the inter-human disagreement with a mean of 10 pixels (median 9 pixels). To put the position error in anatomical perspective, it can be compared to the marmosets’ head size. The width of the head, measured as the inter-tuft-distance, was on average 47 pixels (20 to 80 pixels, varying with the distance to the camera). Thus, the median position error is about 70% of the average head width. In comparison, the inter-human disagreement was 10 pixels, that is, 22% of the average head width. Finally, in some video frames the subjects looked out of the horizontal plane, or head was occluded. Such frames were labeled as “*angle-does-not-apply*” and assigned a 29^th^ class when training the model. An example of this can be seen in the lower panel of **Fig 2B**. During an interval around frames 25 – 50 the test data was labeled with *angle-does-not-apply*, and except for a few frames right at the start and end of the interval predicted by the model as such. Since this was a binary classification we report the error as the area under the receiver operating characteristic curve (AUC). Perfect classification would result in an AUC of 1, and chance classification would result in an AUC of 0.5. The AUC for predicting *angle-does-not-apply* on the complete test set was 0.859.

The errors presented above were the results of averaging over the output of several models. However, the gains made by this procedure were rather limited. The average errors for individual models had a mean of 14.9° (median 7.9°) for direction and 44.8 pixels (median 28.6 pixels) for position. The average *angle-does-not-apply* AUC for individual models was 0.835.

Given information about head direction and position, we investigated where subjects spent most time and what they looked most at (i.e. “point of gaze”). To provide a coarse-grained picture of the most visited locations during an experimental session we calculated two-dimensional histograms of head positions representing the horizontal plane of the test box (see **Fig 3**). Similarly, we estimated the points of gaze through combining direction and position, extrapolating the resulting gaze lines to the perimeter of the test box, and representing them as histograms of looking time. **Fig 3A** shows two examples from an early (upper panel) and a late (lower panel) experimental session, demonstrating how, the subject increasingly directs its attention towards the reward dispenser. This agrees with the fact that, during the late session, the subject spent most time looking at and/or sitting next to the dispenser.

**Fig 3.**
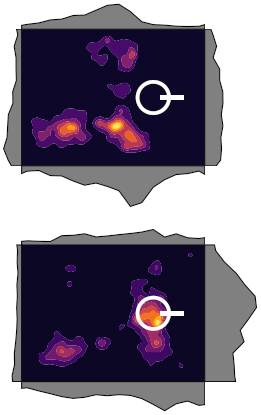
Points of gaze and head positions during two experimental sessions. The density of head positions is shown as heat maps spanning the horizontal plane of the test box. The white horizontal bars and circles on the right shows the locations of the reward dispenser. Along the perimeter of the heat map (i.e. the test box), points of gaze are shown as gray histograms. The upper panel shows an early session before the subject was used to receiving reward from the dispenser while the lower panel shows a late session where the subject’s positions and points of gaze are dominated by the reward dispenser.

However, for a more fine grained analysis of the subject’s behavior during particular times of interest we analyzed how the gaze and position changed frame-by-frame. The video recordings were done during a vocal learning experiment, in which the subjects received a reward two seconds after the end of a contact *phee* call [58]. **Fig 4A** shows three examples of gaze and positions associated with call and reward. Note how the subject looks toward the reward dispenser immediately after the call and then moves to the dispenser. This behavior is further described in **Fig 4B** and **C** where we aligned the animal’s gaze behavior around the time of call and reward. We limited the analysis to the first ten calls since the reward motivation decreased during an experimental session, resulting in less time-locked behavior. The figures demonstrates how the subject directs the gaze towards the dispenser less than a second after the reward is delivered (**Fig 4B**) and soon after start moving towards the dispenser.

**Fig 4.**
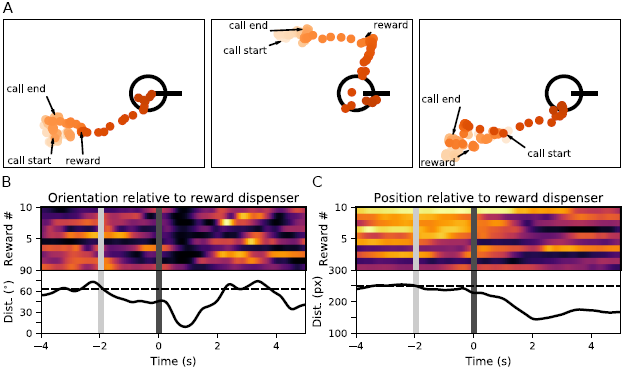
Gaze and head position associated with call and reward. **A**, Examples of gaze direction and head position around the time of reward delivery represented in the horizontal plane of the test box. The gaze and head position is indicated by the orange lines and circles, respectively. The progression of time is indicated by color darkness. For visual clarity, the lines representing gaze not directed at the dispenser are made semi-transparent. Call onset, offset and reward delivery are indicated by arrows, and the black circle and vertical bar to the right of each panel indicate the location of the reward dispenser. **B**, Head direction relative to the reward dispenser from four seconds before to five seconds after reward delivery. The top portion shows the angular distance of the first ten rewards (warmer colors represent greater distance), and the bottom portion shows the average (black trace) with 95% CI (light gray field). The dashed line indicates the baseline (i.e. the average before call offset). The x-axis shows time in seconds, the vertical light gray bar at −2 s indicates the time of call offset, and dark gray bar at 0 s indicates the time of reward delivery. **C**, The spatial distance to the dispenser; otherwise the figure layout follows **B**.

In order to visualize the pattern of movement associated with calls we aligned speed data (i.e. the magnitude of change in direction and position) to the beginning and end of calls. Since the call duration varies between calls, we re-sampled the data during the calls to a common duration (2.67 s, which is the median call duration). This allowed us to align call associated data across multiple calls. **Fig 4** shows how the angular speed (Fig 4A) and head speed (Fig 4A) change around and during a call.

Both the angular speed and the head speed gradually drop by approximately 50% during the calls. This observation is similar to the general decrease in body movement during calls reported by [3]. They also report a slower build-up of speed preceding calls. However, that could not be addressed in the current experiment due to the confounding effect of the reward. As can be seen in **Fig 4A**, the marmosets slow down when consuming the reward, and when finished speed up again. This would create a spurious appearance of a call-associated speed-up.

## Discussion

Here we provided a demonstration of how deep learning can be used to track marmoset gaze behavior. We trained a CNN on video data to predict a marmoset’s head direction and position, giving an estimate of the marmoset’s location and where it was looking. The tracking performance was good, especially for head direction for which the disagreement between the model and human was close to the inter-human disagreement **(Fig 2C)**. For position, the disagreement between human and model was greater, but still comparable to human performance. This level of performance was not unexpected since deep learning algorithms applied to other complex tasks have been shown to perform at [34], or above human levels [19].

Vision is the dominant sensory modality of diurnal primates such as marmosets, making gaze tracking essential to the study of their behavior. Their high acuity and forward facing eyes provides them with excellent binocular vision for fast and accurate guidance in a complex three-dimensional environment [27]. However, measuring gaze behavior is fraught with difficulties. Gaze tracking in non-human primates requires the head to be immobilized, in order to leave eye movements as the sole contributor to gaze shift. Head restraint is generally achieved with an implanted metal head post [33], or in a more recent and less invasive development for macaques, with thermo-plastic helmets [13,31]. Critically, due to the requirement of head restraint, these methods are hard, if not impossible, to apply in several naturalistic settings where gaze tracking would be desirable. In addition, head restraint might have adverse effects on the gaze behavior of primates such as marmosets, where head movements make a big contribution to the gaze shift. To our knowledge, the only alternative to head restrained gaze tracking in non-human primates is to manually score video recordings to get a coarse estimate of gaze direction [9,15]. Beyond being time consuming and thus not feasible to apply to hours of data, the method is very imprecise. In light of these methodological limitations, we believe that the method presented here provides an important tool for future studies of marmoset behavior. However, we would like to reiterate that the method presented here relies on the great contribution of head movements to the gaze shift of marmosets, and is thus not applicable to gaze tracking of species where head-independent eye movements contribute more to the total gaze shift.

The method presented here is flexible and can easily be applied to similar tasks without any changes to the source code (available at https://github.com/kalleknast/head-tracker). To apply it to novel video recordings of isolated marmosets, one would merely need to label a subset of the sdata and train the model. We annotated a total of 20,127 frames which took around 20 hours. However, a subset of those (5,000 frames) was used to select model architecture and to evaluate performance. Thus, 15,000 frames (15 hours of work) are sufficient to train the model. Since annotating video is the main time-sink when applying the model to new data, it is worth to consider how to further minimize this effort. One possibility would be to selectively label the more informative frames. Video frames are often highly correlated, meaning that many of them are redundant, and thus do not contribute much when optimizing the model. Through the selective sampling of less correlated frames the model could probably be successfully trained using less labeled data [51]. It is thus likely that not more than 10 hours needs to be spent on data labeling in order to apply the method presented here to a novel set of video recordings.

Although our experiments were performed with isolated individuals, we believe that the general method can be extended to recordings of multiple animals. Such simultaneous tracking of multiple animals would provide a way to quantify social behaviors, which are in general, complex and difficult to measure, typically requiring sophisticated analysis of high dimensional data such as video. In the past, social behaviors could only be measured by manually scoring videos, making this type of analysis low-throughput [46], inconsistent, and subject to unconscious and/or conscious bias [8]. However, deep learning based methods, such as the one presented here, hold the promise of automating this task. Marmosets, like humans, look directly at regions of social interest, particularly faces [33]. Thus, by tracking their gaze in a social environment, we get a measure of their social interactions. Gaze and position tracking would provide the time series data that could be used for automatic detection of such behaviors as play and grooming. The method presented here provides a first step in that direction.

The recordings were performed in a restricted environment that essentially only enabled two-dimensional movement. The tracking was further constrained to only concern directions and positions in the horizontal plane, thus decreasing the complexity of our implementation. However, marmosets are not ground dwelling, and in most naturalistic environments tracking in three dimensions is necessary for a meaningful description of their activities. While three-dimensional tracking requires multiple cameras synced by a common clock, the tracking algorithm would not require much modification. The simplest way would be to train independent two dimensional models on video from different cameras, and to reconstruct three-dimensional coordinates and angles *post hoc*. Successful, multi-camera, three-dimensional tracking has been reported for *Drosophila* [52,53]. However, that method relies on background subtraction to localize the target, and thus, requires a visually constant and unobstructed environment which precludes extension to primate home-cage conditions. The utilization of deep learning algorithms provides an alternative that is much more robust to the visually messy environments of marmoset cages.

A model’s performance is constrained by the quality of the input data, which in our case was limited by the precision of the direction labels (i.e. the width of the direction bins) and the accuracy of the human-scored training data. However, in spite of these limitations of the input data, the model’s performance was very close to the human level. Since the error introduced by data binning is randomly distributed, it should be reduced by prediction averaging. For example, averaging over 30 models resulted in a direction error of 10.9°, whereas the average error of individual models was 14.9°, significantly greater. Similarly, the error of individual models could also be decreased (from 14.9° to 13.7°) by computing the predicted direction as the sum over labels weighted by the softmax activation. That is, treating the softmax output as a probability distribution over the labels. However, none of these strategies are likely to reduce the error caused by the uncertainty in the human assigned labels, thus leaving this as the lowest error a model can achieve. With this in mind the direction error should be considered close to optimal.

The error in position was relatively greater, and most likely caused by the loss of precision due to the down-sampling of the video frames by a factor of 16 (from 640 × 480 to 160 × 120 pixels). That is, an error of one pixel in the model’s output translates to an error of 16 pixels in the full sized video frame. This limit is inherent to the design of the current model, and is probably best dealt with through building an explicit object detection model.

## Materials and Methods

### Dataset Description

The subjects were three captive-born adult common marmosets (two females and one male), housed at the Instituto do Cérebro, Universidade Federal do Rio Grande do Norte. The marmosets were housed socially in two wire mesh enclosures (1.20 × 1.50 × 2.45 m), enriched with tree branches, ropes, plants, hammocks and nesting tubes. The animals were fed twice daily with fresh and dried fruit, nuts, egg and chicken, and had *ad libitum* access to water. The colony was maintained outdoors protected by a roof while still allowing daily sunbaths in natural light. The animals were housed in compliance with SISBIO permit #18394, and the experiment was approved by the ethics committee of Universidade Federal do Rio Grande do Norte with CEUA permit #028/2013. No animal was sacrificed at the end of the experiment.

We recorded video of marmosets taking part in a vocal learning experiment. During an experimental session, an individual marmoset was kept acoustically isolated in a test box measuring approximately 30 × 30 × 30 cm **(Fig 1)** for 15 minutes. The subjects were trained to voluntarily leave their home-cage and enter the test box cage in return for preferred food items. The subjects were rewarded for making contact *phee* calls [58]. Sound was monitored online via a microphone streaming data to a computer running a *phee*-detection program [59]. Two seconds after the end of a *phee* call, a dispenser delivered a liquid reward (Yakult, Tokyo, Japan) via a spout to a reward recipient placed below (**Fig 1** and **2A**). We recorded 70, 15-min videos (total of 17 hours and 30 min) over a period of six months. A total of 4002 *phee* calls were recorded. For a subset of the experimental sessions (12 videos from one subject), the recordings allowed for precise alignment of the video frames and the behavioral events. Video was recorded at 15 frames per second with a C920 Logitech HD Pro Webcam (Logitech, Lausanne, Switzerland) placed 30 cm above the cage. The recording software was custom written in Python 3.5 utilizing the GStreamer open-source multimedia framework (https://gstreamer.freedesktop.org) and is available at https://github.com/kalleknast/video-tools.

Subsequent to the data collection, a subset of the video frames were annotated with the marmosets’ head position and orientation in the horizontal plane. In order to reduce effort and improve consistency, we wrote software to aid the annotation process. Annotation did not require any previous training since it only consisted of marking the location of the marmoset’s forehead followed by the base of the two tufts. From these three points, the head direction and position in the horizontal plane was calculated. However, in some instances, frames could not be assigned any meaningful head direction in the horizontal plane. For example, sometimes the subject looked out of the horizontal plane (i.e. up or down), or its head was obscured. Such frames were annotated with *angle-does-not-apply*. We annotated 18,127 frames from 15 different videos (1208 ±413 frames per video). These frames were split into a training set (15,127 frames) and a validation set (3,000 frames). The validation set was used to tune the model architecture (e.g. to find the optimal number of classes, see below). Another 2,000 frames were annotated for the test data. Test data was taken from five videos, each contributing 400 frames that were sampled in four randomly spaced batches of 100 consecutive frames. Since the temporal correlation between consecutive video frames is high using randomly drawn frames from the same videos as used for training data would probably overestimate the model’s ability to generalize to new data. Thus, in order fairly assess the model’s performance, the videos used for test data were different from those used for validation and training data.

Video annotation was done by two investigators. In order to estimate their agreement and to get a reference to compare the model’s performance to, both investigators annotated the same 1,361 video frames. There was a good agreement about both head direction (mean 9.2° and median 7.2°, 0 to 180° range), and position (mean 10 pixels and median 9 pixels, 640 × 480-pixel frames). It took approximately one hour to label 1,000 video frames (3.6 s per frame), and around 20 h to annotate the complete dataset.

### Classification Algorithm

Our goal was to find a model able to predict both the head’s direction and its position in the horizontal plane. In order to achieve this, we took advantage of the fact that the units of a CNN have spatially restricted receptive fields [63], and thus, implicitly carry information about the location of the source of their activation. That is, the units act as object localizers despite not receiving any explicit information about the location of the object in the image. However, most CNN architectures incorporate one or more layers of fully-connected units in the final stages, thus losing the position information. Here, we instead followed [65] and replaced the fully-connected layer with a global average pooling layer, averaging over filter features while retaining spatial information [26,65]. This layer was, in turn, connected to a softmax classification layer. In order to find the discriminative image regions, the softmax output was mapped back to the last convolutional layer resulting in a class activity map (CAM) [65] that, for head localization, was up-sampled from 1024 units to 160 × 120, that is, to the same shape as the input.

Following this strategy we built a CNN with six convolutional layers and two max pool layers. Filter sizes in the convolutional layers decreased from 11 × 11 to 3 × 3. The global average pooling had a size of 1024 and connected to a softmax layer with the number of units equal to the number of classes. We used cross-entropy loss regularized by the Euclidean (L2) norm of the weights the regularization parameter set to 0.0001. Weights were initialized with random values drawn from a normal distribution with zero mean and a standard deviation set to 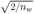 where *n_w_* is the number of weights in a layer [19]. The model was optimized with the Adam algorithm [26], and was trained for 80 epochs. Inputs to the model were video frames, whitened, down-sampled (from 640 × 480 to 160 × 120 pixels) and converted to gray-scale (averaged RGB color). Over-fitting is a common problem for models built with a great number of tunable parameters. We addressed this in two ways: through regularization (see above), and model averaging [50,51].

Head direction measured in degree of head rotation in the horizontal plane was binned in *n* – 1 classes and the frames labeled as *angle-does-not-apply* were assigned to an *n^th^* class. In order to select the optimal number of classes, and thus, the size of direction bins, we trained multiple models configured to predict from five to 33 classes (steps of four), and evaluated their performance on the direction error on the validation set. The error decreased with an increasing number of classes until *n* = 29, where it reached the inter-human disagreement. Based on this, the final model was trained to predict head direction angles converted into 28 classes (0 to 360° in 28 bins, each ≈ 12.9° wide) and a 29^th^ class representing *angle-does-not-apply*. The head position was approximated as the location of maximum activity of the up-sampled CAM (see **Fig 2A**).

We used Python 3.5 to implement the head tracking algorithms. The core model was written using GPU-enabled TensorFlow 0.10.0 [1], while NumPy SciPy [60] and Matplotlib [22] were used for additional functionality. The software is available at https://github.com/kalleknast/head-tracker. Computations were done on a Lenovo ThinkStation P900 running Ubuntu 16.10 with a 4 GB NVIDIA Quadro K2200 graphics card.

### Data Analysis

We evaluated the model’s performance on both direction and position. The position error was measured as the distance in pixels between the human-assigned position and the coordinates of peak activity of the CAM, up-sampled to the same size as the model’s input. However, the direction error had to be treated in two separate parts. First, for *angle-does-not-apply* (i.e. presence/absence of a discernible head direction in the horizontal plane), we report the area under the ROC-curve (AUC). This metric was chosen instead of standard accuracy since the *angle-does-not-apply* label only occurs in approximately 12% of the frames (i.e. the classes were unbalanced). Second, we report the error in predicted direction as the angular distance between the human-assigned label and the model’s prediction converted back from 28 classes to degrees of angle. **Fig 2B** demonstrates examples of the direction and position errors. We report both the average errors of individual models, and the errors of all models combined (i.e. the result of model averaging).

To investigate the subjects’ points of gazes during the experimental sessions, we combined the head direction and position through extrapolating a ray beginning at the estimated head position and having an angle given by the head direction. We computed histograms of the end points of these “gaze-rays” along the perimeter of the test box (see **Fig 3A**). Position preference was estimated by computing two-dimensional histograms representing the density of head positions in the horizontal plane of the test box (see **Fig 3A**). For visual clarity, the density map was smoothed with a Gaussian kernel having a 10 × 10-pixel standard deviation.

Reward-associated behavior was analyzed by extracting data in an interval starting four seconds before to five seconds after the rewards. We limited the analysis to the first 10 rewards delivered within an experimental sessions since the number of calls, and thus rewards, varied between experimental sessions, and the subjects’ interest in the rewards tended to decrease during a session. From this reward-aligned data we calculated the distance in pixels between the head position and reward dispenser, and the angular distance between the gaze-ray dispenser (i.e. how many degrees the head would need to be rotated in order to be directed towards the dispenser). The time points where the model predicted *angle-does-not-apply* were interpolated from neighboring values. For both measures, we smoothed the 10 traces with a Gaussian kernel (13 ms SD) and computed averages and 95% confidence intervals (CI) (see **Fig 4B** and **C**).

The movement speeds associated with the production of the contact *phee* calls were analyzed similarly. We extracted the angular speed of head rotation (the magnitude of change in head direction), as well as, the head speed (the magnitude of change in head position) from five seconds before to 2.5 seconds after the calls. However, since the calls varied in duration we re-sampled the data from the start to the end of the call to the median call duration (2.67 s), allowing us to align the data across calls (see **Fig 5**).

**Fig 5.**
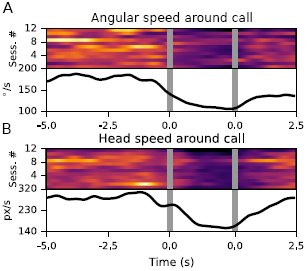
Speed around the time of calls. **A**, Angular speed of head rotation. The upper panel show the average angular speed around calls per session (warmer colors indicate greater speed). The lower panel show the angular speed averaged over sessions (black) ±95% CI (gray field). Call start and end is indicated by the two vertical grey bars. x-axis show time in seconds. **B**, Head speed; otherwise the figure layout follows **A**.

## Acknowledgments

We thank Stephen Shepherd and Daniel Y. Takahashi for comments on the manuscript, and William Wood, Josy Pontes, Janaina Figueiredo, Bruno Soares, Maria Bernardete de Souza and the Primatology Center of UFRN for help with various aspects of housing & specialized marmoset care. Research equipment and HKT’s and TBRC’s stipends were funded by CNPq/BJT grant # 402422/2012-0 to SR and HKT. SR further thanks the FAPESP Research, Innovation and Dissemination Center for Neuromathematics (grant # 2013/07699-0, S. Paulo Research Foundation). We thank Taeksoo Kim for guidance on the implementation of the CNN model.

